# Noisy information about the environment: A source of individual differences within and across generations

**DOI:** 10.64898/2026.05.15.725332

**Authors:** Nicole Walasek, Marjolein Bruijning, Karthik Panchanathan, Willem E. Frankenhuis

**Affiliations:** Evolutionary and Population Biology, Institute for Biodiversity and Ecosystem Dynamics, University of Amsterdam, Amsterdam, the Netherlands; Department of Anthropology, University of Missouri, USA; Max Planck Institute for the Study of Crime, Security and Law, Freiburg, Germany

**Keywords:** individual differences, stochastic variation, non-shared environment, cue sampling, phenotypic plasticity

## Abstract

Despite sharing the same genes and the same environment, individuals often develop substantial phenotypic differences. While this pattern has been documented across diverse species and traits, the processes giving rise to this “stochastic” or non-shared environmental variation remain unclear. Recent mathematical models of development in which phenotypes are gradually constructed may offer some clues. These models show that stochastic sampling of imperfect environmental cues can generate striking variation in developmental trajectories and adult phenotypes. At the population level, such imperfect cues produce increasing stability of individual differences across ontogeny (e.g. animal personality) and patterned distributions of mature phenotypes (e.g. normal or skewed) that resemble those observed in real organisms. Our paper synthesizes existing models in which stochastic phenotypic variation arises solely as a by-product of mechanisms missing their phenotypic targets because of imperfect cues. We then link these models to related, but independent, mathematical theory exploring the environmental conditions under which stochastic phenotypic variation is favoured by natural selection. Our integration shows that stochastic sampling is often favoured over classic bet-hedging strategies involving non-plastic generalist or specialist strategies. Our findings provide new directions of research on stochastic sampling as a mechanism for adaptive stochastic variation within and across generations.

## Introduction

> *“No one supposes that all the individuals of the same species are cast in the very same mould. These individual differences are of the highest importance for us […] and they thus afford materials for natural selection to accumulate.” (from Charles Darwin, 1859, the Origin of Species)*

Phenotypic variation is the raw material from which evolution by natural selection forges adaptation. Biologists typically focus on three sources of variation: genetic variation among individuals, the induction of phenotypic variation by environmental variation, and interactions between genetic and environmental sources of variation. Yet, a growing body of empirical work documents large and consistent phenotypic differences among genetically identical individuals living in the same environment [1,2]. This residual variation—henceforth *stochastic variation*—persists even after controlling for genetic and shared environmental influences [2] and is widespread across organisms, from insects and fish, to rodents and humans [3–8]. For example, behavioural geneticists have long acknowledged non-shared environmental influences as a major driver of phenotypic variance among children growing up in the same family [9]. Despite this mounting evidence for stochastic variation across species and traits, most studies still prioritize explaining and tracking changes in trait means rather than trait variation. Here, we centre the variation by exploring and linking mechanisms of stochastic variation that operate across both developmental and evolutionary timescales.

### The development of stochastic variation

Multiple mechanisms generate stochastic variation, some operating within a generation and others across generations. Within a generation, developmental noise from factors including gene expression, cell differentiation, and epigenetic regulation, can result in phenotypic variation among genetically identical individuals raised in controlled environments [10–12]. It is less well known that stochastic variation can result even in the absence of such noise when developmental trajectories are influenced by the sampling of environmental information.

Adaptive phenotypic plasticity, in which organisms adjust traits to environmental conditions [13,14], requires organisms to infer the environmental state through the acquisition and integration of environmental cues. This information acquisition process is inherently noisy as environmental cues are imperfectly correlated with environmental states, a process we call ‘stochastic sampling’ [15,16]. For example, two identical organisms estimating predation risk from occasional predator sightings may, by chance, observe different numbers of predators and thus infer different levels of predation risk.

A growing body of models conceptualize adaptive phenotypic plasticity as a sequential decision making process, in which organisms gradually learn about and adapt to their environment across development using imperfect cues [17–20]. These models of incremental development show that stochastic sampling across ontogeny can, by itself, generate substantial phenotypic variation among genetically identical individuals living in the same environment [15,21–27]. Crucially, this variation is often structured rather than idiosyncratic. Models of incremental development produce consistent among-individual differences across ontogeny, individual differences in plasticity, and characteristic distributions of mature phenotypes that dovetail with empirical patterns [28–31]. This research highlights the central role played by an organism’s *information state* as an intermediary linking genotype and environment to phenotype. Even when genetically identical individuals experience the same environment, stochasticity in sampling can cause their estimates of the environment to diverge. These differences can, in turn, cause differences in developmental trajectories. From this perspective, individual differences may emerge as a by-product of organisms attempting to optimally match their phenotypes to their environments using imperfect information.

### The evolution of stochastic phenotypic variation

While stochastic phenotypic variation may arise merely as a by-product of development, it may also be a trait under selection. Stochastic variation is defined as consistent phenotypic differences among genetically identical individuals experiencing the same environment. Following Lewontin’s criteria [32], stochastic variation can be a target of selection if genotypes within a population differ in their production of stochastic variation, these differences are heritable, and they result in fitness differences. Evidence relevant to the first criterion comes from clonal systems, in which genetically identical individuals develop phenotypic differences even when living in the same environment [2,4,5,33]. Crucially, comparisons among genotypes show that the extent of this variation differs within populations. For instance, inbred lines of fruit flies differ in their maze-navigation behaviour [3,34] and isogenic strains of mice show large differences in within-strain behavioural variability [35].

The second criterion, heritability, is supported by several lines of evidence. The most direct evidence comes from genome-wide association studies in fruit flies, where the degree of variability among individuals has been shown to be heritable and associated with specific loci [3,36]. Artificial selection experiments show that variance itself can also respond to selection in mammals. In mice, lines have been successfully bred to differ in variability of birth weight [37]. Heritable trait variation resulting from artificial selection is often modest, ranging around ∼0–5% of the total variance [38], but can sometimes be as high as 21% (e.g. for birth weight in mice [37]). Across species, new statistical methods can track how both the mean and variance of a trait evolve over evolutionary time. These methods show, for example, that beak length varies more among individuals in forest-living bird species compared to species occupying other habitats [39]. Heritable non-genetic mechanisms, such as epigenetic parental effects, can also contribute to stochastic variation, but such effects will not be our focus here [40].

Lastly, research indicates that stochastic variation can impact fitness. Stochastic variation is expected to be favoured whenever increased trait variability enhances long-term fitness, for instance through bet-hedging or nonlinear averaging. Bet-hedging is an evolutionary strategy that acts on phenotypic variation among offspring to reduce variance in fitness across generations [41–43]. When environments fluctuate unpredictably and phenotypic optima cannot be anticipated, producing a wide range of phenotypes can increase long-term geometric mean fitness across generations, at the expense of arithmetic mean fitness within a generation. Bet-hedging may take the form of conservative bet-hedging, in which individuals adopt intermediate phenotypes that achieve moderate fitness across a range of environments; or diversified bet-hedging, in which genotypes produce different (specialised) phenotypes, each of which performs well in some environments but poorly in others. Empirical support for bet-hedging includes classic examples of variable seed germination timing that buffer lineages against environmental unpredictability [44]. Nonlinear relationships between trait values and fitness provide a complementary mechanism for producing adaptive stochastic variation. When the trait–fitness relationship is convex, Jensen’s inequality implies that a distribution of varied phenotypes yields higher mean fitness than the mean phenotype alone [45]. Variability resulting from such nonlinear averaging may therefore be adaptive even in stable environments, as shown for instance by the interaction between the mean and variance of plant toxin concentrations and their effects on herbivores [46].

### Bridging timescales and perspectives

Theoretical research on stochastic variation has developed along two complementary but largely independent lines: models on the evolution and development of phenotypic plasticity explain how imperfect cues generate consistent individual differences within lifetimes, while evolutionary models explore when phenotypic variation is favoured across generations through mechanisms such as bet-hedging and nonlinear averaging. Bridging these perspectives raises a key question: Can stochastic sampling—a within-generation consequence of organisms using imperfect environmental cues— produce phenotypic variation that is also adaptive across generations?

We address this question by synthesising results from a family of existing mathematical models [47] on the evolution and development of phenotypic plasticity [24,25,27]. These models assume that organisms have identical genotypes and experience the same environment. Organisms sample imperfect cues across development to reduce uncertainty about the environmental state and incrementally adjust their phenotypes to match environment states. Within each model, we vary the long-term distribution of environmental states to which organisms have adapted (the ‘evolutionary prior’) and the degree to which cues reduce environmental uncertainty (the ‘cue reliability’). For any combination of evolutionary prior and cue reliability, individual differences arise purely from stochastic sampling. Aggregating results across these different combinations allows us to disentangle the genetic, environmental, and information processing contributions to stochastic variation. We then compare the evolutionary success of the model-derived phenotypic distributions, expressed as long-term geometric mean fitness, to non-plastic specialist and generalist strategies.

Our analyses address two goals. First, we systematically map how genes, environments, and information processing shape stochastic variation. Second, we ask whether stochastic sampling within generations can produce stochastic variation that is adaptive across generations.

## Methods

Model code and data can be found at https://osf.io/x8g6k/overview?view_only=150107f74a7e49d8b7a4ab337bb9ef65.

### General model of incremental specialisation

At the onset of ontogeny, the period relevant for the development of a trait, organisms randomly disperse into discrete and non-overlapping patches. Each patch can be in one of two environmental states (*E*_0_ or *E*_l_), and we initially assume that the environmental state remains constant across ontogeny. The organism does not know the state of its patch but is adapted to the long-term distribution of environmental patches. The organism uses this distribution as an evolutionary prior estimate of being in either of the two states denoted by *P*(*E*_0_) and *P*(*E*_1_) [48]. The more this prior deviates from a uniform distribution (*P*(*E*_0_) = *P*(*E*_1_) = 0.5), the more likely one state is than the other at the onset of ontogeny.

Ontogeny lasts for 10 discrete time periods. In each period, an organism samples a cost-free environmental cue (*C*_0_ or *C*_1_) to learn about the state of its patch. Cues are imperfect indicators of the underlying environmental state. The cue reliability specifies the conditional probability of observing the correct cue in the corresponding state (*P*(*C*_0_|*E*_0_) and *P*(*C*_1_|*E*_1_)). We assume that *P*(*C*_0_ |*E*_0_ ) = *P*(*C*_1_ |*E*_1_ ) and that the probability of observing the wrong cue corresponds to *P*(*C*_1_|*E*_0_) = 1 − *P*(*C*_0_|*E*_0_) and *P*(*C*_0_|*E*_1_) = 1 − *P*(*C*_1_|*E*_1_). The higher the cue reliability, the better cues help to reduce uncertainty about the environmental state. Organisms use Bayesian inference to optimally integrate information obtained from cues with their priors [16,48,49].

Organisms incrementally tailor their phenotypes to the underlying environmental state across ontogeny. We assume that there is an optimal phenotype for each environmental state (*P*_0_ and *P*_1_ ), which correspond to the fully specialised phenotypes for *E*_0_ and *E*_1_. These two phenotypic targets (*P*_0_ and *P*_1_) should be conceptualized as independent trait dimensions rather than two ends of a continuum (e.g. ornaments to attract mates and defensive morphology). As a result, adjustments to one trait do not automatically affect the other. However, there is an implicit trade-off between the two traits resulting from opportunity costs: time invested in one leaves less time for investing in the other.

Specifically, in each period an organism samples a cue and then makes one of three phenotypic decisions: (1) develop one incremental specialisation towards *P*_0_, (2) develop one incremental specialisation towards *P*_1_, or (3) wait and forgo specialisation. While sampling environmental cues and adjusting phenotypes are cost-free, there is nevertheless a trade-off: Organisms that start specialising early in ontogeny and stay on target will end up with a higher level of phenotypic specialisation compared to organisms that specialise and then switch based on new information.

Organisms accrue fitness based on how well their phenotype at maturity—the time window after ontogeny has ended—matches the environmental state. Fitness increases based on the number of correct incremental adjustments, and decreases based on the number of incorrect adjustments. We explore three mappings between phenotype and fitness rewards and penalties: marginally linear, marginally increasing, and marginally diminishing [24,25,27]. When fitness functions are increasing, many phenotypic adjustments are needed to reap large fitness benefits. When fitness functions are diminishing, few phenotypic adjustments can have large fitness benefits.

For each combination of prior and cue reliability, we use stochastic dynamic programming with backwards induction to compute optimal developmental policies [49–51]. An optimal policy specifies the fitness-maximising decision in every possible state across ontogeny. An organism’s state at any time period *t* consists of their current phenotype and cues sampled (for equations, see [24,25,27]).

### Plasticity across ontogeny

Plasticity at a given time in ontogeny is the extent to which subsequent phenotypic development depends on cues. For each model and all possible parameter combinations, we now simulate pairs of cloned individuals who follow the same optimal policy, experience the same sequences of cues, and thus make the same phenotypic decisions. These clones develop together until time period *t* after which they are separated. From then on, one clone receives the opposite cues compared to its counterpart until the end of ontogeny. When one clone samples *C*_0_, the counterpart samples *C*_1_ and vice versa. Plasticity then corresponds to the Euclidean distance between mature phenotypes of paired clones. We normalize this measure to range between 0 and 1 by dividing by the maximally attainable Euclidean distance between the moment of separation and the end of ontogeny. For each pair of clones, we then compute the average plasticity level across ontogeny.

### Model variants

1. ***Stable cue reliability, stable environment***. The baseline model assumes that both the cue reliability and the environmental state are constant across ontogeny [27].
2. ***Varying cue reliability, stable environment***. In the first variant of the baseline model, the environmental state is constant across ontogeny, but the cue reliability varies between 0.55 and 0.95 [24]. We explore three patterns of change between these extremes: (1) cue reliability continuously increases, (2) cue reliability first increases then decreases (triangular), and (3) cue reliability continuously decreases. All three patterns result in the same average cue reliability of 0.75. At any time *t*, organisms update their estimates of the environmental state using the current cue reliabilities *P*(*C*_0,*t*_|*E*_0_) and *P*(*C*_1,*t*_|*E*_1_).
3. ***Stable cue reliability, varying environment***. In the second variant, the cue reliability is fixed, but the environmental state varies between *E*_0_ and *E*_1_ across ontogeny [25]. The transition probabilities *P*(*E*_1_|*E*_0_) and *P*(*E*_0_|*E*_1_) determine the rate of change, yielding autocorrelations of 0.2, 0.5, or 0.8. The relationship between autocorrelation values *r* and transition probabilities is: *r* = 1 (*P*(*E*_1_|*E*_0_) *P*(*E*_0_ |*E*_1_)). Higher autocorrelation values imply more stable environments. Transitions can be symmetric (both directions equally likely) or asymmetric (one direction more likely). The long-term state distribution (the evolutionary prior *P*(*E*_0_), *P*(*E*_1_)) follows from these transition probabilities. Symmetric transitions yield *P*(*E*_0_) = *P*(*E*_1_) = 0.5. Asymmetric transitions bias the long-term distribution towards one state. In this model, organisms accrue fitness in adulthood under a fluctuating environment. Fitness corresponds to the average match between the mature phenotype and the environmental state across adulthood. We vary the ratio of adulthood to ontogeny by considering a short adulthood (1 adult period) and a long adulthood (20 periods), while fixing ontogeny to 10 periods, thus capturing variation in the duration of the juvenile period relative to maturity (a life-history trait).

### Quantifying stochastic variation

Figure 1 summarizes the workflow from model specification to stochastic variation. For combinations of evolutionary prior, cue reliability, and, where applicable, pattern of cue reliability change, rate of environmental change, and duration of adulthood, we compute three measures: (1) variation in mature phenotypes, (2) average rank switches across ontogeny (as a measure for variation in developmental trajectories), and (3) individual differences in plasticity. In the main text we only describe and present results for the first measure. We provide descriptions of and results for the other measures, which yield qualitatively similar results, in the supplements.

**Figure 1:**
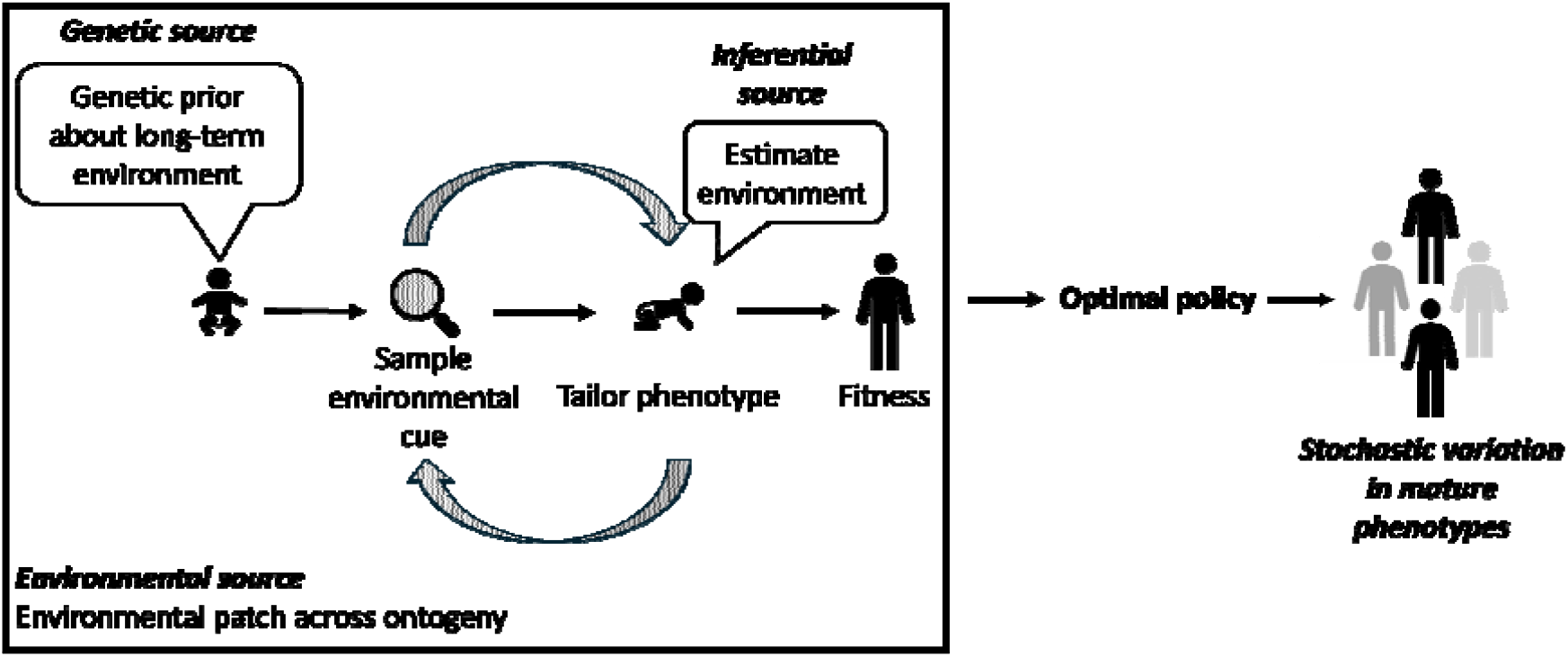
General model of incremental specialisation. Organisms start ontogeny with an evolutionary prior over the long-term distribution of environmental states. In each time period organisms sample a cost-free environmental cue and update their estimates of the environment based on that cue. Organisms then incrementally tailor their phenotypes to the expected long-term environment. This cycle continues until the end of ontogeny when organisms accrue fitness. Model variant (1) (the baseline model) assumes stable cue reliability and environmental state within generations. Variants (2) and (3) relax these assumptions by introducing within- and across-generation variation in cue reliability and in environmental state, respectively (both indicated by striped filling). Stochastic dynamic programming then allows us to identify optimal developmental policies for different combinations of priors, cue reliabilities, and rates of environmental change. Based on the optimal policy we then simulate populations of developing organisms and, finally, quantify stochastic variation in distributions of mature phenotypes. The shading darkness of simulated individuals indicates different levels of specialisation towards one phenotypic trait.

For each model, we quantify stochastic variation by using the optimal policy to simulate developmental trajectories and distributions of mature phenotypes across all possible parameter combinations (see Figure 1). Phenotypes consist of three numbers: the number of time periods spent specialising towards *P*_0_, the number of time periods spent specialising towards *P*_1_, and the number time periods spent waiting (foregoing specialisation).

To quantify variation in mature phenotypes, we focus on one of the two phenotypic traits. For models assuming a stable environmental state within generations, we pick the trait that corresponds to the actual underlying state. For the model assuming changes in the environmental state across generations, we pick the trait corresponding to the more likely state. We then create a histogram of that trait for each simulated population of mature individuals. We normalize trait values to range between 0 and 1 by dividing them by the highest trait value possible (10). If the organism spent all 10 time periods specialising towards the trait, the normalized value is 1. From the histogram we compute a single measure of phenotypic variation defined as the product of two normalized quantities: (1) the standard deviation of trait values, capturing the spread in trait values; and (2) the Shannon information entropy of the trait-value distribution, capturing how evenly spread trait values are. Both components are normalized to range from 0 to 1 by dividing by their theoretical maxima. The resulting product equals 1 when each possible trait value occurs with equal probability, and 0 when the entire population has the same trait value.

### Geometric mean fitness of optimal policies and fixed strategies

We compare the geometric mean fitness of optimal policies adapted to stable environments within generations to policies adapted to changing environments within generations, as well as with two fixed strategies. To this end, we include the two models that only differ in the presence of within-generational environmental changes (i.e. the baseline model [27] and the second variant [25]). For both models, we use the optimal policy to simulate distributions of mature phenotypes across all possible parameter combinations. We simulate populations developing (or, in case of within-generation environmental changes, starting to develop) in *E*_0_ and *E*_1_. Additionally, we compute distributions of mature phenotypes resulting from two distinct fixed strategies: generalists and specialists. Generalists produce phenotypes that contain mixtures of specialisations towards *E*_0_ and *E*_1_. The relative contribution of the two types of possible specialisations is proportional to the prior. Specialists produce fully specialised phenotypes (i.e. fully specialised towards *E*_0_ or *E*_1_), where the proportion of the two possible specialists is proportional to the prior distribution.

For each population resulting from optimal policies and fixed strategies, we then compute fitness (i.e. *F*_0_ and *F*_1_) in each environmental state as the sum of fitness rewards for correct and penalties for incorrect specialisations (see above). We assume that, across generations, the environment alternates between the two states based on the prior distribution. Long-term fitness, calculated as the geometric mean fitness, of a population, then corresponds to: . We add the resulting geometric mean to a baseline fitness constant to ensure positive fitness values across generations.

## Results

### Synthesis of models: stochastic variation as a by-product of developmental processes

Figure 2 shows stochastic variation quantified as the phenotypic variation in mature phenotypes. We show plots for other measures, which yield qualitatively similar results, in the supplements (Figures S1; S4-S7). Across all models, the more prior uncertainty there is about the long-term environmental state (i.e. the long-term distribution over environmental states is closer to 0.5), the greater the amount of phenotypic variation that results from stochastic sampling. Thus, the ‘genetic’ source, representing long-term adaptations to the distribution of environmental states, influences the overall amount of stochastic variation that will emerge. In addition, we observe effects of the environmental source. Stochastic sampling from more heterogenous experiences across ontogeny (i.e. lower cue reliability across or early in ontogeny), results in larger magnitudes of stochastic variation when the environmental state is stable (Figure 2, top row). The degree of heterogeneity in experiences is less related to the magnitude of stochastic variation when the environmental state changes across ontogeny (Figure 2, bottom row). Additionally, we observe larger magnitudes of stochastic variation when adulthood is short. Thus, stochastic sampling interacts with genetic and environmental sources, as well as life-history traits, to produce varying degrees of stochastic variation. Lastly, we observe that high levels of plasticity do not always produce large magnitudes of stochastic variation while low levels do not exclude the development of stochastic variation (see Figure S6). Thus, levels of plasticity alone do not determine how much stochastic variation emerges.

**Figure 2:**
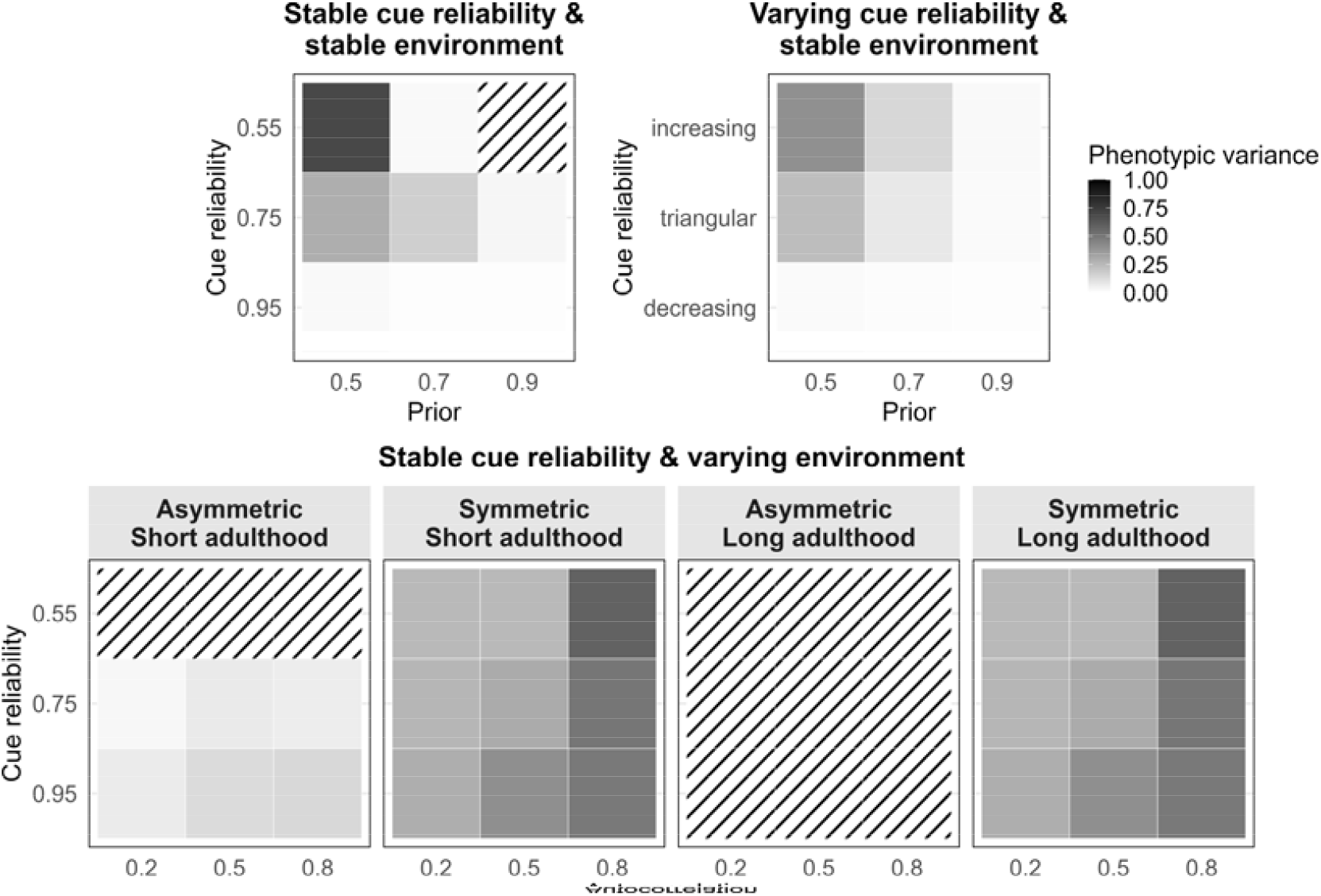
Phenotypic variation in mature phenotypes (linear fitness rewards and penalties). The top left panel depicts a model with stable cue reliability and stable environmental state across ontogeny. The top right panel indicates a model with varying cue reliability and stable environmental state across ontogeny. The horizontal x-axes indicate the prior estimate of one environmental state and the vertical y-axes the cue reliability (top left) and pattern of cue reliability changes (top right). The bottom row depicts a model with stable cue reliability and varying environmental state across ontogeny. The two leftmost panels depict results for a short adulthood and the two rightmost panels for a long adulthood. For a given duration of adulthood, transition probabilities can be asymmetric, meaning that one environmental state is more likely than the other in the long term, or symmetric, meaning that both states are equally likely. Within each panel the horizontal x-axis indicates the autocorrelation and the vertical y-axis the cue reliability. Tiles with striped diagonal lines indicate parameter combinations that result in zero plasticity across ontogeny. The z-axis (colour scale) is scaled to range between the measure’s theoretical minimum and maximum.

When the environment is stable, patterns of phenotypic variance, rank switches, and individual differences in plasticity are qualitatively similar across linear, increasing, and diminishing rewards (Figures S4–S15, top row). In changing environments, however, the shape of fitness rewards matters (Figures S4–S15, bottom rows). With increasing rewards, phenotypic variance is largely independent of the rate of environmental change and higher for low and moderate cue reliability (Figure S8). Compared to linear rewards, increasing rewards favour more specialised phenotypes, resulting in distributions resembling diversified bet-hedging across generations. We also observe fewer rank switches and individual differences in plasticity overall, suggesting that cues have less impact across ontogeny (Figures S9-S11). With diminishing rewards, lower rates of environmental change and higher cue reliability result in more phenotypic variance (Figure S12). When organisms have little information, they tend to develop generalist phenotypes to reap some of the diminishing rewards, resulting in less phenotypic variance and distributions resembling conservative bet-hedging across generations. Because organisms switch specialisation trajectories more often, diminishing rewards generate more rank switches and individual differences in plasticity across combinations of rates of environmental change and cue reliability (Figures S13-S15). In conclusion, the shape of the reward function substantially affects stochastic variation when environments change across ontogeny, but not when they are stable.

### When is phenotypic variation from stochastic sampling favoured across generations?

The preceding analyses show how stochastic sampling generates phenotypic variation as individuals use imperfect cues to optimally tailor their phenotypes to their environments. We next ask whether the phenotypic distributions produced by these optimal developmental policies are also favoured across generations.

To address this question, we compare long-term fitness from our optimal policies to two distinct fixed strategies: a generalist strategy expressing an intermediate trait value, and a specialist strategy expressing two specialised trait values (Figure 3). We note that under linear rewards and penalties, specialists and generalists have identical fitness: because long-term fitness scales linearly with specialisation, distributing specialisation across individuals (specialists) or within individuals (generalists) in proportion to the prior yields the same population fitness (supplements Table S1-S2). This equivalence breaks down under nonlinear fitness functions. When fitness is increasing, concentrating specialisation in a subset of individuals (specialists) can be favoured; when fitness is diminishing, spreading specialisation within individuals (generalists) can be favoured (supplements Table S1-S2). This is consistent with the broader principle that nonlinear averaging governs the adaptiveness of stochastic variation: the shape of the fitness function can determine both whether stochastic variation is favoured and what form it takes.

**Figure 3:**
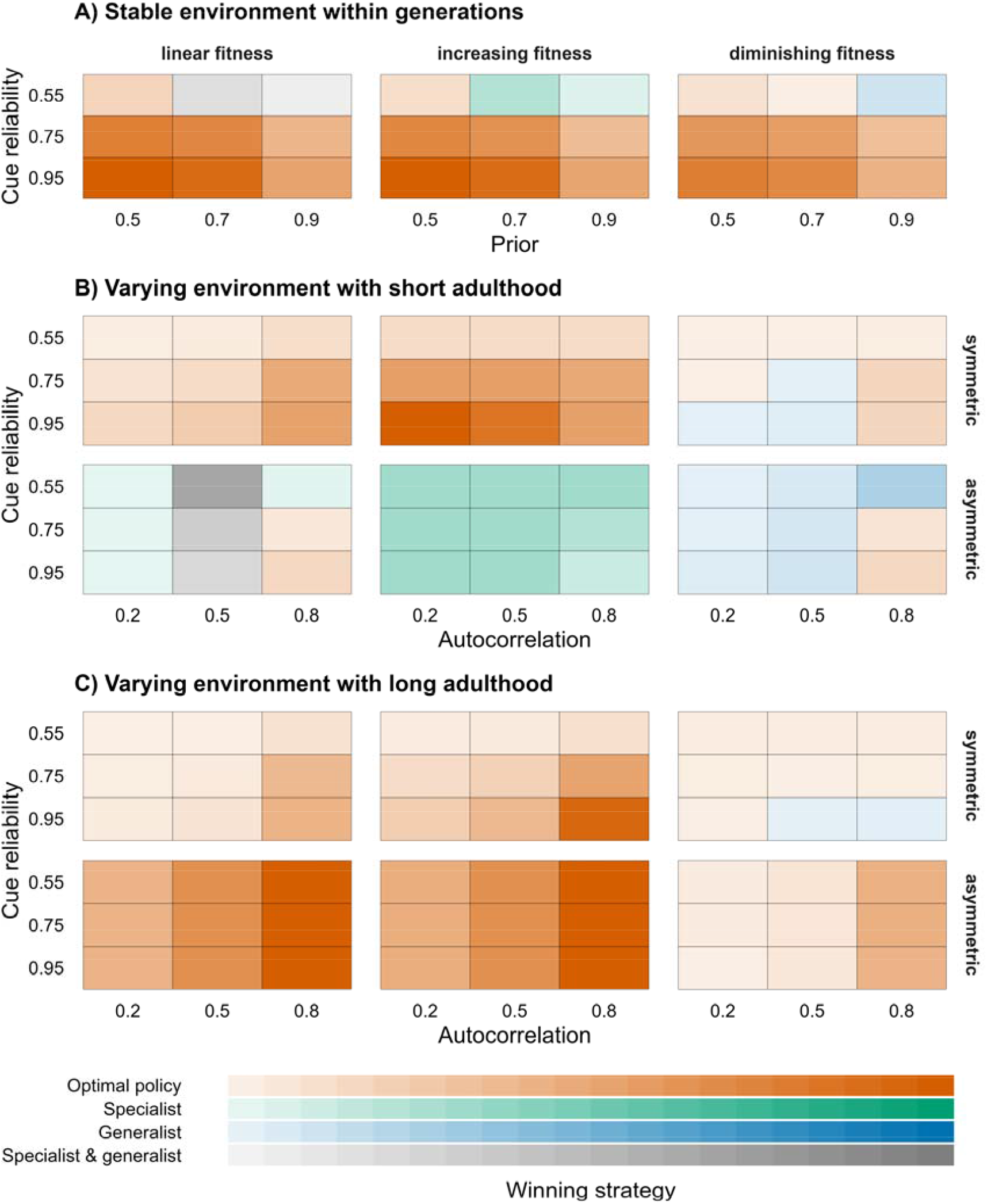
Strategies that yield the highest geometric mean fitness. For different combinations of prior, cue reliability, autocorrelation, and duration of adulthood, we compute geometric mean fitness across generations with the phenotypic distributions that emerge from our optimal policies (allowing for plasticity). We also compute geometric mean fitness of a fixed (non-plastic) specialist and a generalist strategy. Across all panels (A-C), columns indicate different fitness reward functions. Panel A corresponds to a model assuming an environment that varies across generations, but that is stable within generations. Within each subpanel the horizontal x-axis depicts the prior estimate of one environmental state. The vertical y-axis depicts the cue reliability. Panels B and C correspond to a model assuming an environment that varies both within and across generations. Rows represent either symmetric (uniform prior distribution) or asymmetric (prior distribution biased towards one state) transition probabilities between environmental states. Within each subpanel the horizontal x-axis depicts the autocorrelation of environmental changes. The vertical y-axis depicts the cue reliability. Colours indicate the winning strategies which achieved the highest geometric mean fitness, with grey indicating a tie between the generalist and specialist fitness values. The intensity of the colour indicates the difference in magnitude between the winning and second best strategy. Exact fitness values are shown in the supplements (Tables S1-S2).

Our analysis reveals that optimal policies often outperform fixed strategies (orange shading in Figure 3), especially when cues are more reliable and environments stable within a generation (panel A). When the environmental state can change within generations, fixed strategies can outperform the optimal policy, particularly when adulthood is short and one environmental state is more likely than the other (panel B, bottom row). Under these conditions, the optimal policy tends to produce phenotypes specialised towards the more likely environment. While this strategy fares well in the majority of generations, it can produce substantial mismatch across generations featuring the rare environment. When adulthood is short, a single timestep can determine fitness within a generation, leading to potentially large mismatch if it happens to be the rare environment. Thus, across generations, environmental conditions during these short adulthoods are effectively more unpredictable, favouring non-plastic generalist or specialist strategies.

Whenever conditions favour a fixed strategy (a specialist or generalist), the strategy qualifies as bet-hedging: here, the fixed strategy results in higher geometric mean fitness than the optimal policy but lower arithmetic mean fitness within a generation, evidenced by the fact that the optimal policy differs from the fixed strategy [52]. The specialist strategy corresponds to diversified bet-hedging, whereas the generalist strategy corresponds to conservative bet-hedging. In line with previous work on nonlinear averaging, increasing fitness rewards favour diversified bet-hedging while linear or decreasing rewards favour conservative bet-hedging [53]. More broadly, our findings align with work showing that bet-hedging is favoured in unpredictably changing environments [54]. In contrast, when adulthood is long, optimal policies tend to outperform fixed strategies by leveraging prior information and cues (or sometimes by ignoring cues in non-plastic policies), to tailor phenotypes to the long-term environment (panel C).

Thus, our analysis highlights that developmental mechanisms that optimally use cues to tailor phenotypes can themselves produce adaptive stochastic variation to cope with environmental fluctuations across generations. Moreover, stochastic variation arising from such optimal development is most likely to be selected for when organisms have access to reliable cues in stable or slowly changing environments. In unpredictably changing environments, we should expect other mechanisms, such as non-plastic bet-hedging strategies, to play a larger role in generating adaptive stochastic variation. Together, these first explorations suggest that stochastic variation emerging as a by-product of stochastic sampling can also be adaptive across generations.

## Discussion

In this paper, we integrated findings from optimality models of gradual development of individual differences. These models share the assumption that stochastic phenotypic variation arises as a by-product of developmental mechanisms missing their phenotypic targets due to imperfect cues. Our results show that individuals with the same genetic prior in the same environment often diverge because they encounter different sequences of environmental cues. These sequences generate different estimates of environmental conditions, which in turn result in phenotypic variation. The contribution of such stochastic sampling to phenotypic variation is largest when there is prior uncertainty about the long-term environment and when cues have low reliability, producing heterogenous cue sets. We further show that these effects interact with within-generational rates of environmental change and lifespan. Moreover, we compared long-term population fitness achieved by stochastic sampling to two fixed strategies (generalists and specialists) and found that stochastic sampling is often favoured. Aggregating across these results, we see that uncertainty about the long-term environmental conditions often, but not always, favours an optimal policy resulting in high levels of phenotypic variation across generations.

Our analyses highlight stochastic sampling as a potential mechanism for producing adaptive stochastic variation, underscoring the importance of organisms’ information states [55]. Our results show that variation in cues sampled within the same environment alone can generate substantial individual differences. Our modelling further shows that the shape and amount of individual differences arising from informational processes can be predicted from features of the developmental system (genetic priors) and the information structure of the environment (cue reliability). This process differs from, yet is compatible, with other developmental sources of phenotypic variation, such as stochasticity in molecular and (epi)genetic processes [10–12]. Stochastic sampling additionally complements work on ‘luck’ as a driver of fitness differences among otherwise identical individuals [56,57]. This work emphasizes how random sequences of environmental events (‘environmental stochasticity’) generate variation in demographic and fitness outcomes. Our results suggest that this part of variation may be implemented via differences in organisms’ information states arising from stochastic sampling.

### Stochastic sampling as an adaptive mechanism operating on developmental timescales

Building on these general results and given the formal similarities between adaptive processes operating on different timescales [58,59], it is worth considering adaptations to environmental unpredictability on shorter timescales as well. In our models, environmental unpredictability sets the parameters of developmental mechanisms, which produce stochastic variation within generations. Do developmental experiences, in turn, shape the parameters of mechanisms for adjusting to unpredictability on shorter timescales, such as during moment-to-moment interactions?

This question dovetails with research in cognitive neuroscience on the ‘hyperparameters’ that control learning processes [60]. The idea is that people set general learning parameters—such as the balance between exploration and exploitation—based on their developmental histories, and then fine-tune parameter values to specific contexts. For example, a child might initially avoid exploration in a new school, because it has learned to be cautious at home, but increase exploration after learning the school is safe. Strikingly, children start learning about the degree of unpredictability as early as infancy [61]. Particularly relevant here is the ‘adaptive stochasticity hypothesis’. This hypothesis proposes that developmental exposure to unpredictability upregulates the production of stochastic neurocognitive phenotypes to better cope with short-term environmental change [62]. Such phenotypes may arise from randomness in transcription and translation of critical proteins, axonal outgrowth, and spontaneous neuronal activity (i.e. developmental noise). The authors reason that in predictable environments, stochasticity may be costly because it risks producing maladaptive phenotypes; yet in unpredictable environments, stochasticity can provide flexibility and robustness. Consistent with their hypothesis, the authors show that children from low socioeconomic backgrounds—who experience more unpredictability, on average—exhibit more stochasticity in neurodevelopment than children from high socioeconomic backgrounds.

We propose that stochastic sampling of information in unpredictable environments can drive the development of these observed phenotypic differences. Our results may also help explain why some children show greater neurodevelopmental stochasticity than others despite growing up in similarly unpredictable environments [63,64]: variation in information states generated by stochastic sampling could amplify or attenuate the impact of environmental unpredictability. Future work can formally investigate whether stochastic sampling favours adaptive stochasticity by exploring when developmental exposure to unpredictability should increase sampling-driven variation over variation from other developmental mechanisms. Predictions derived from these models can then be compared against empirical patterns of neurodevelopment and behaviour.

### Stochastic sampling as a mechanism for bet-hedging

At an evolutionary timescale, previous work has shown that bet-hedging can be an adaptive evolutionary strategy to cope with environmental unpredictability [41–43,54], but the mechanisms that generate bet-hedging remain poorly understood [44]. We propose the novel hypothesis that stochastic sampling accompanying developmental plasticity can serve as such a mechanism. Prior work has often contrasted plasticity with bet-hedging rather than viewing developmental plasticity as a mechanism that effectively leads to bet-hedging [54]. In our models, plastic (and non-plastic) optimal policies frequently outperform fixed, non-plastic bet-hedging strategies. Under some environmental conditions, our plastic policies produce phenotypic distributions that resemble those obtained from fixed bet-hedging strategies. But, unlike fixed bet-hedging strategies, the optimal policy uses available information to fine-tune distributions to the experienced environmental state, resulting in a form of plastic bet-hedging. Strictly speaking, these policies maximise both arithmetic and geometric mean fitness given available information and so do not qualify as bet-hedging, which requires a reduction in arithmetic mean fitness. Yet, empirical work supports the idea that bet-hedging can be fine-tuned through plasticity [65,66], and one theoretical model shows that plasticity can reduce fitness variance across generations under conditions typically thought to select for bet-hedging [67].

Future work using evolutionary agent-based simulations [68] could examine the evolution of plastic bet-hedging, in which optimal policies tailor individuals to their environments both within and across generations while maximising long-term (geometric mean) fitness (see [51] for a discussion of why dynamic programming is not feasible for such questions). Beyond bet-hedging of phenotypic specialisation, such models could extend prior modelling work [22] by investigating bet-hedging of differential plasticity, that is, the co-existence of plastic and non-plastic strategies. Taken together, such work may clarify when plasticity fine-tunes bet-hedging and whether this process itself generates variation in plasticity across individuals.

In conclusion, our analyses identify stochastic sampling as a mechanism for adaptive phenotypic variation. By using imperfect environmental information to guide development, organisms can produce stochastic variation that is adaptive both within lifetimes and across generations.

