## supplements for "Noisy information about the environment: A source of individual differences within and across generations"

### Computing other forms of stochastic variation

#### *Average rank switches across ontogeny*

As with the variance in mature phenotypes, we pick one trait to follow to measure variation in developmental trajectories. At each time period, we normalize trait values to range between 0 and 1. We then rank individuals based on their trait value in each time period. The more specialized individuals are towards the trait of interest, the higher their rank in that time period. Organisms with the same trait value share a rank. Between two consecutive time periods we compute the proportion of individuals that experienced a rank switch. Our final measure corresponds to the average proportion of rank switches across ontogeny, thus providing a summary of overall variation in developmental trajectories. Additionally, this measure allows us to approximate between-individual variances in a trait across ontogeny. Such between-individual variances constitute the main source of consistent individual differences in animal personality (i.e. repeatability) [41,42].

#### *Individual differences in plasticity*

Plasticity at a given time in ontogeny is the extent to which subsequent phenotype development depends on cues. For each model and all possible parameter combinations, we now simulate pairs of cloned individuals who follow the same optimal policy, experience the same sequences of cues, and thus make the same phenotypic decisions. These clones develop together until time period $t$ after which they are separated. From here on, one clone receives the opposite cues compared to their counterpart until the end of ontogeny. That is, whenever one clone sees $C_0$, the counterpart sees $C_1$ and vice versa. Plasticity then corresponds to the Euclidean distance between mature phenotypes of paired clones. Additionally, we normalize this measure to range between 0 and 1 by dividing by the maximally attainable Euclidean distance between the moment of separation and the end of ontogeny. For each pair of clones, we then compute the average plasticity level across ontogeny. Finally, we compute the standard deviation in plasticity levels across all simulated pairs of clones as a measure of individual differences in plasticity.

### Averages across all stochastic variation measures

#### *Stochastic variation—linear fitness rewards & linear penalties*

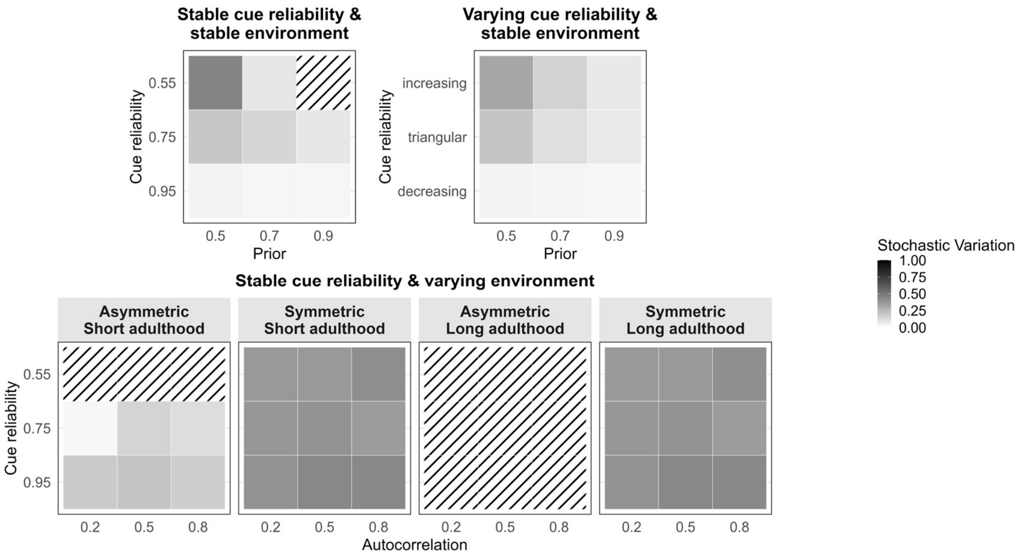


*Figure S1:* Averages across stochastic variation measures. Stochastic variation corresponds to the unweighted average across variation in mature phenotypes, average rank switches, and individual differences in plasticity. Fitness rewards and penalties are both linear. The top left panel depicts a model with stable cue reliability and stable environmental state across ontogeny. The x-axis indicates the prior estimate of one environmental state and y-axis the cue reliability. The top right panel indicates a model with varying cue reliability and stable environmental states across ontogeny. The x-axis indicates the prior estimate of one environmental state and the y-axis the patterns of changes in cue reliability. The bottom row depicts a model with stable cue reliability and varying environmental state across ontogeny. The first two panels in the bottom row depict results for a short adulthood and the last two panels for a long adulthood. For a given duration of adulthood, transition probabilities can be asymmetric, meaning that one environmental state is more likely than the other in the long-term, or symmetric, meaning that both states are equally likely. Within each panel the x-axis indicates the autocorrelation and y-axis the cue reliability. Tiles with striped diagonal lines indicate parameter combinations that result in zero plasticity across ontogeny. The z-axis (colour scale) is scaled to range between the measure's theoretical minimum and maximum.

***Stochastic variation—increasing fitness rewards & linear penalties***

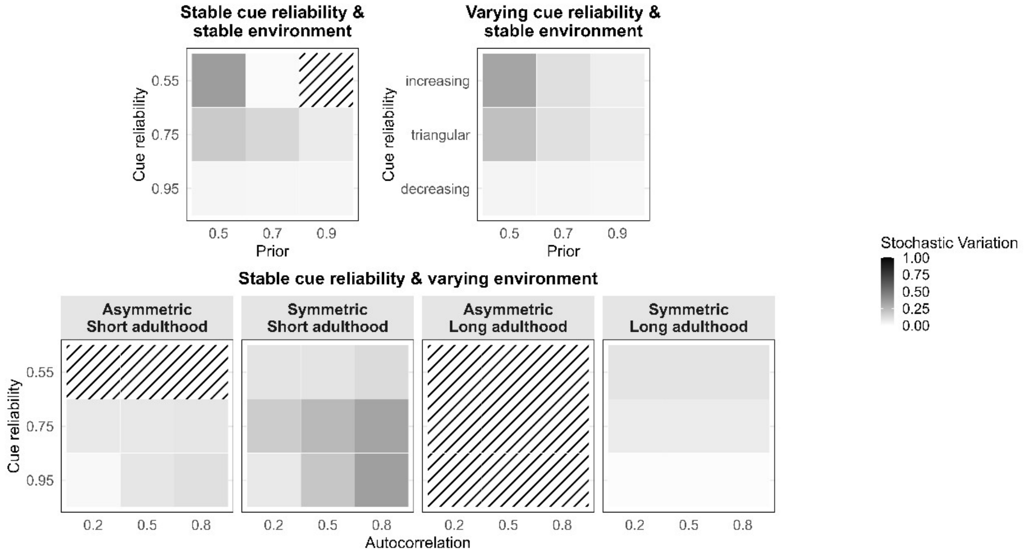


*Figure S2:* Averages across stochastic variation measures. Stochastic variation corresponds to the unweighted average across variation in mature phenotypes, average rank switches, and individual differences in plasticity. Fitness rewards are increasing and penalties are linear. The top left panel depicts a model with stable cue reliability and stable environmental state across ontogeny. The x-axis indicates the prior estimate of one environmental state and y-axis the cue reliability. The top right panel indicates a model with varying cue reliability and stable environmental states across ontogeny. The x-axis indicates the prior estimate of one environmental state and the y-axis the patterns of changes in cue reliability. The bottom row depicts a model with stable cue reliability and varying environmental state across ontogeny. The first two panels in the bottom row depict results for a short adulthood and the last two panels for a long adulthood. For a given duration of adulthood, transition probabilities can be asymmetric, meaning that one environmental state is more likely than the other in the long-term, or symmetric, meaning that both states are equally likely. Within each panel the x-axis indicates the autocorrelation and y-axis the cue reliability. Tiles with striped diagonal lines indicate parameter combinations that result in zero plasticity across ontogeny. The z-axis (colour scale) is scaled to range between the measure's theoretical minimum and maximum.

***Stochastic variation—diminishing fitness rewards & linear penalties***

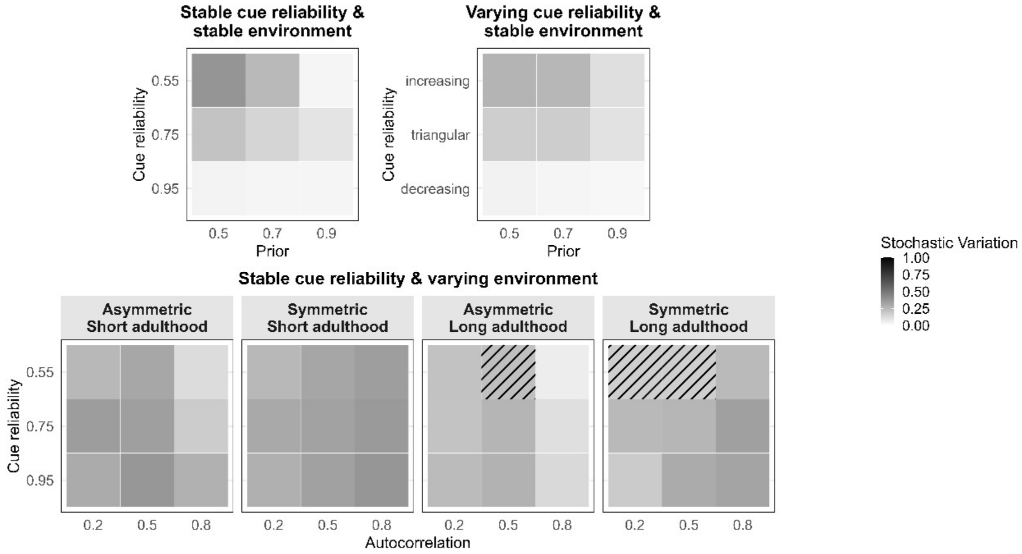


*Figure S3:* Averages across stochastic variation measures. Stochastic variation corresponds to the unweighted average across variation in mature phenotypes, average rank switches, and individual differences in plasticity. Fitness rewards are diminishing and penalties are linear. The top left panel depicts a model with stable cue reliability and stable environmental state across ontogeny. The x-axis indicates the prior estimate of one environmental state and y-axis the cue reliability. The top right panel indicates a model with varying cue reliability and stable environmental states across ontogeny. The x-axis indicates the prior estimate of one environmental state and the y-axis the patterns of changes in cue reliability. The bottom row depicts a model with stable cue reliability and varying environmental state across ontogeny. The first two panels in the bottom row depict results for a short adulthood and the last two panels for a long adulthood. For a given duration of adulthood, transition probabilities can be asymmetric, meaning that one environmental state is more likely than the other in the long-term, or symmetric, meaning that both states are equally likely. Within each panel the x-axis indicates the autocorrelation and y-axis the cue reliability. Tiles with striped diagonal lines indicate parameter combinations that result in zero plasticity across ontogeny. The z-axis (colour scale) is scaled to range between the measure’s theoretical minimum and maximum.

***Individual measures—linear rewards & linear penalties***

***Variance in mature phenotypes***

Higher variance in mature phenotypes indicates a more evenly spread and wider distribution of mature phenotypes with respect to one trait (Figure S4). Across all models, phenotypic variance is highest when there is prior uncertainty (prior of 0.5 or symmetric transition probabilities) about the long-term environmental state. The lower the rate of environmental change and the lower the reliability of cues, the higher phenotypic variance is. When priors are informative (priors larger than 0.5 and asymmetric transition probabilities) and cue reliability is high, phenotypic variance depends on whether the environmental state can change. In contrast to a stable environmental state, changing conditions result in more phenotypic variance even when cues are highly reliable.

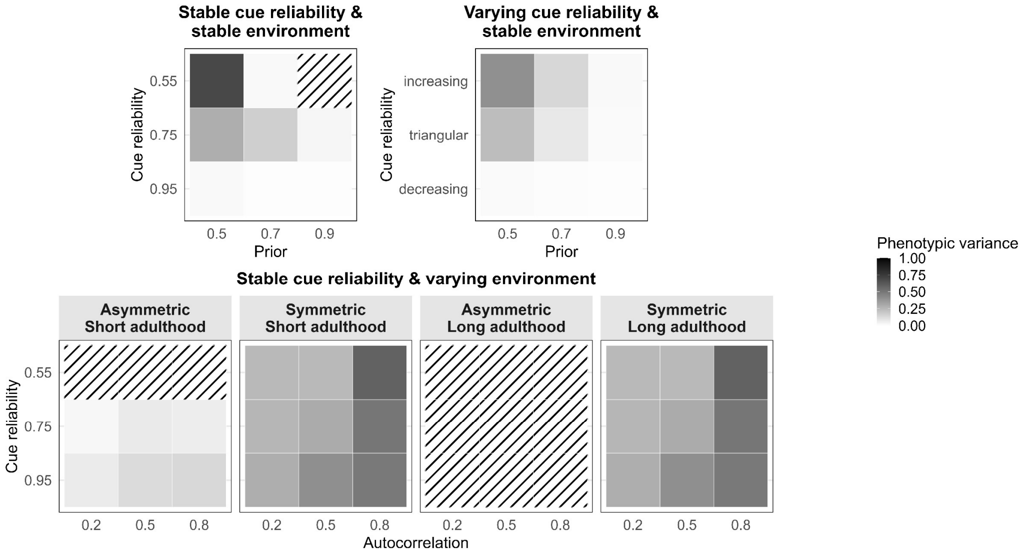


*Figure S4*: Variance in one trait. See, Figure S1 for the remainder of the legend.

***Average rank switches across ontogeny***

The higher average rank switches across ontogeny, the more between-individual phenotypic variability there is across ontogeny (Figure S5). Across all models, rank switches are most frequent when there is prior uncertainty about the long-term environmental state. Under these conditions, higher rates of environmental changes, result in more rank switches. When prior uncertainty is paired with a stable or slowly changing environmental state, lower and moderate cue reliability result in more rank switches. When priors are informative and cue reliability is high, rank switches depend on whether the environmental state can change and on the duration of adulthood. In contrast to a stable environmental state, changing conditions and shorter adulthood result in more rank switches across ontogeny even when cues are highly reliable.

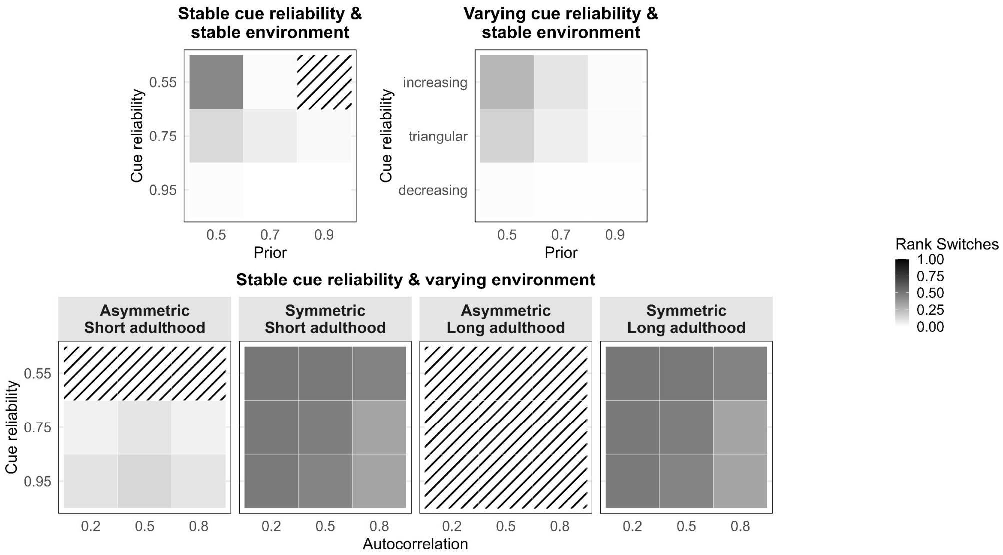


*Figure S5*: Average rank switches. See, Figure S1 for the remainder of the legend.

***Average levels and individual differences in plasticity***

We show average levels of plasticity across ontogeny to contextualize patterns of individual differences in plasticity (Figure S6). When plasticity is non-zero, the level of plasticity is relatively homogenous across models and parameter combinations. We observe the lowest levels of plasticity in a stable environment, either when the cue reliability is low or decreasing. We observe the highest levels of plasticity in a slowly changing environment with highly reliable cues. Compared to average plasticity levels, individual differences in plasticity tend to vary more across models and parameter combinations (Figure S7). Higher individual differences in plasticity indicate higher between-individual variance in average levels of plasticity across ontogeny. When the environmental state is stable, results are qualitatively similar for stable and changing cue reliabilities. Here, individual differences in plasticity are largely independent of prior uncertainty. Additionally, individual differences in plasticity are lowest when cue reliability is highest. When the environmental state can change and plasticity is non-zero, individual differences in plasticity are relatively constant across symmetric and asymmetric transitions probabilities, durations of adulthood, as well as combinations of rates of environmental changes and cue reliability. Here, individual differences in plasticity are lowest, when both states are equally likely (i.e. with uncertain priors) and cue reliability is low or moderate. Notably, the overall magnitude of individual differences is larger when the environmental state changes, compared to models assuming a stable environmental state.

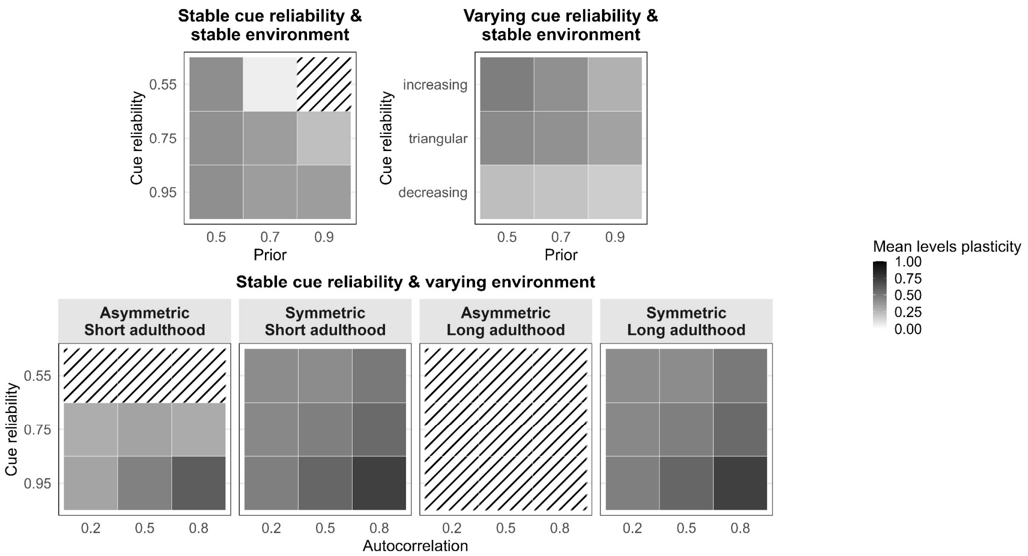


*Figure S6*: Average levels in plasticity. See, Figure S1 for the remainder of the legend.

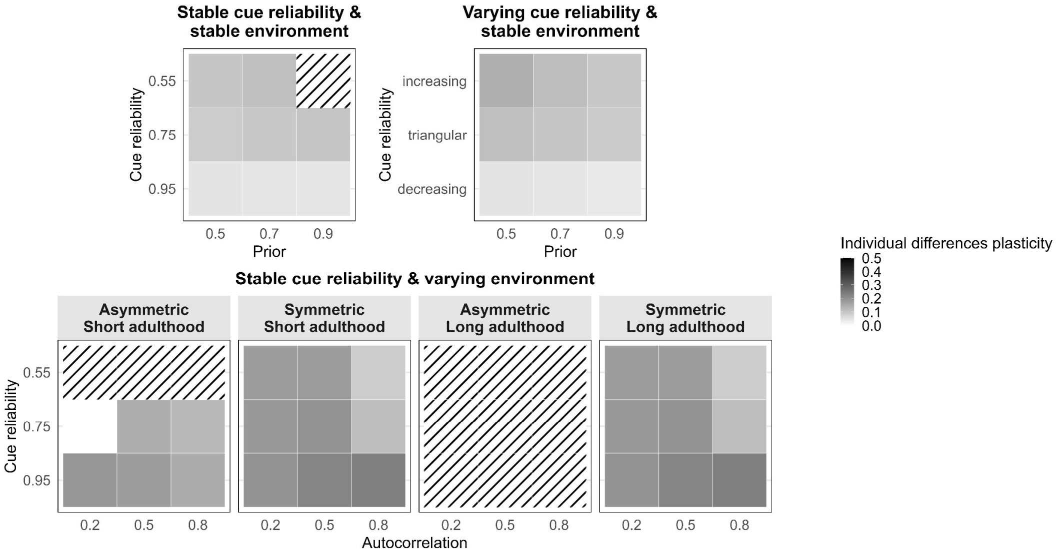


*Figure S7*: Individual differences in average levels of plasticity. See, Figure S1 for the remainder of the legend.

### ***Individual measures—increasing rewards & linear penalties***

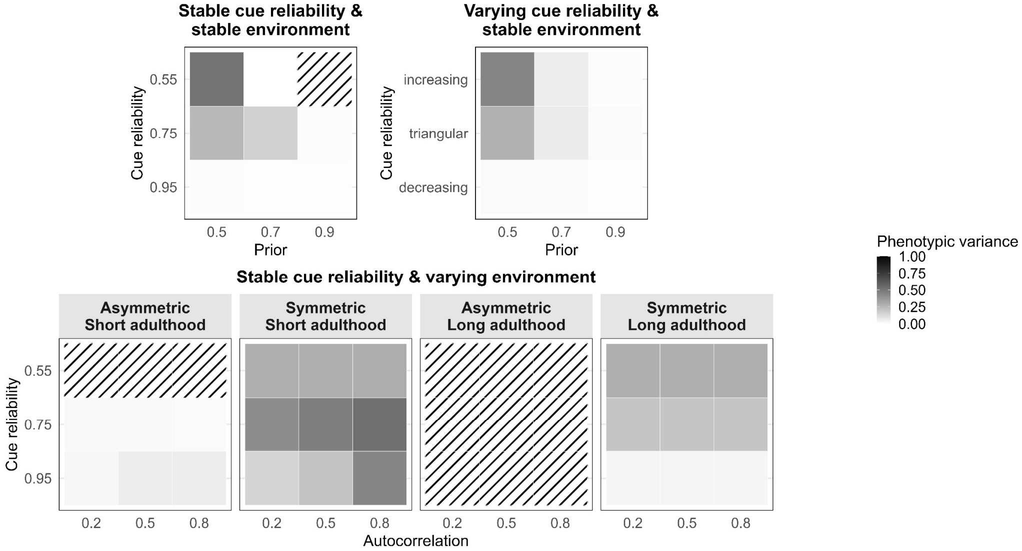


*Figure S8*: Variance in one trait. See, Figure S2 for the remainder of the legend.

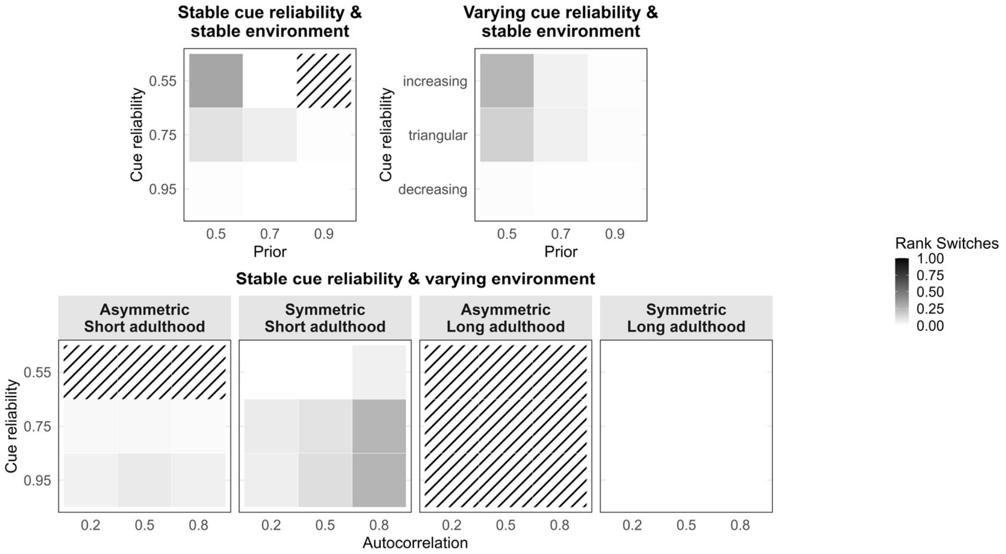


*Figure S9*: Average rank switches. See, Figure S2 for the remainder of the legend.

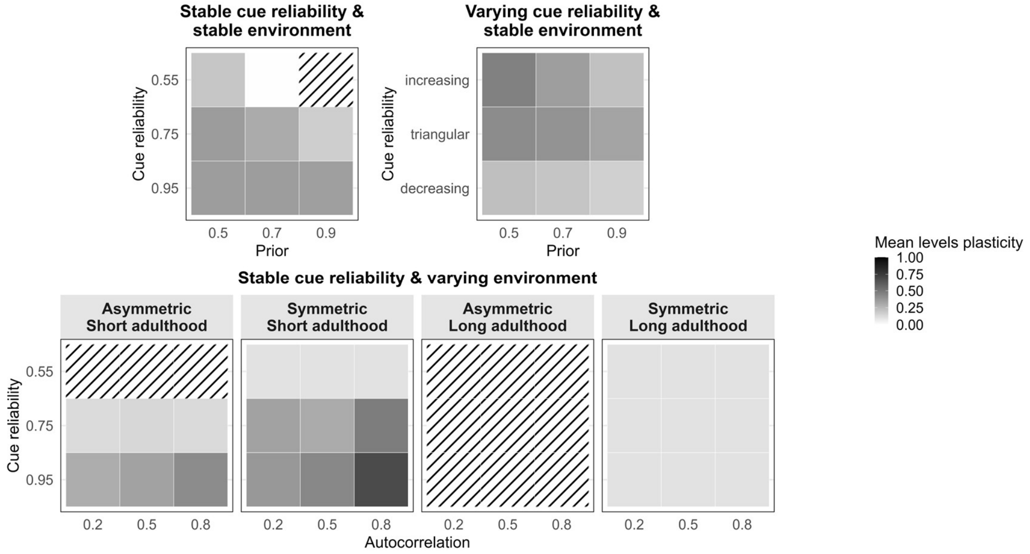


*Figure S10*: Average levels in plasticity. See, Figure S2 for the remainder of the legend.

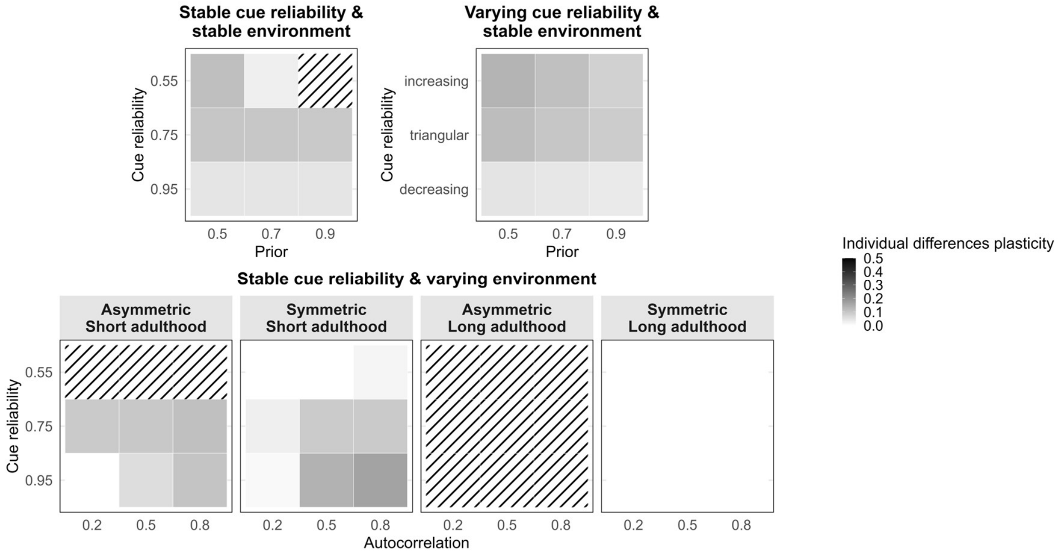


*Figure S11*: Individual differences in average levels of plasticity. See, Figure S2 for the remainder of the legend.

***Individual measures—diminishing rewards & linear penalties***

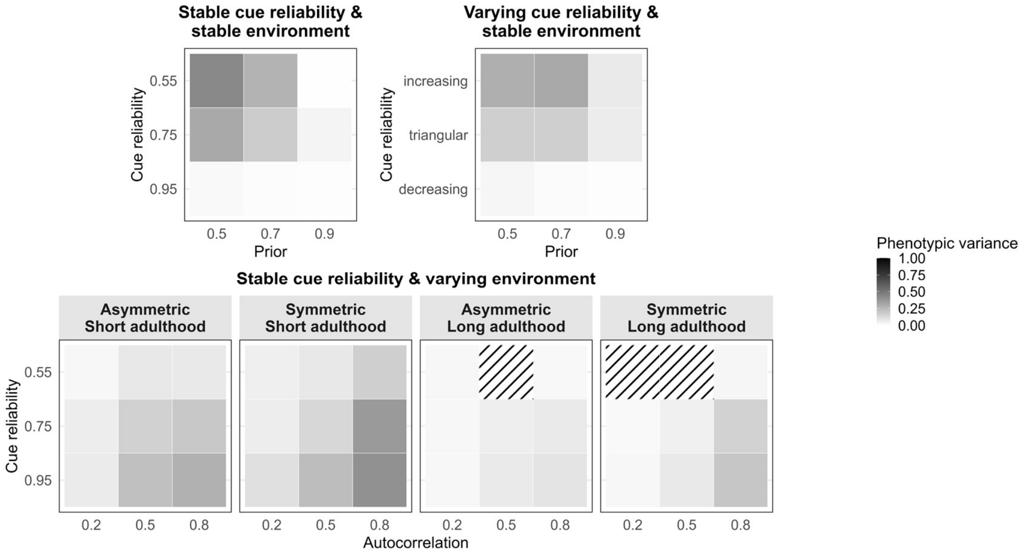


*Figure S12*: Variance in one trait. See, Figure S3 for the remainder of the legend.

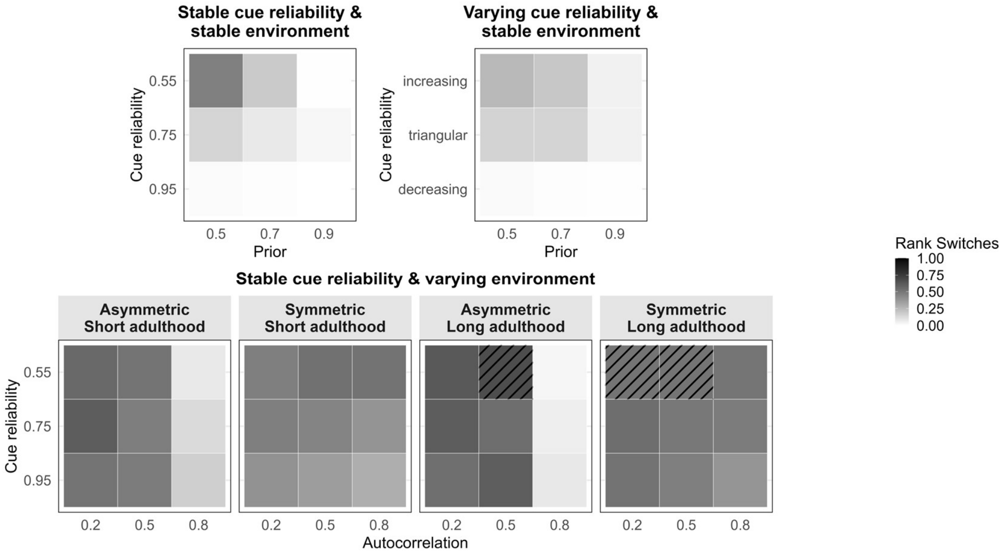


*Figure S13*: Average rank switches. See, Figure S3 for the remainder of the legend.

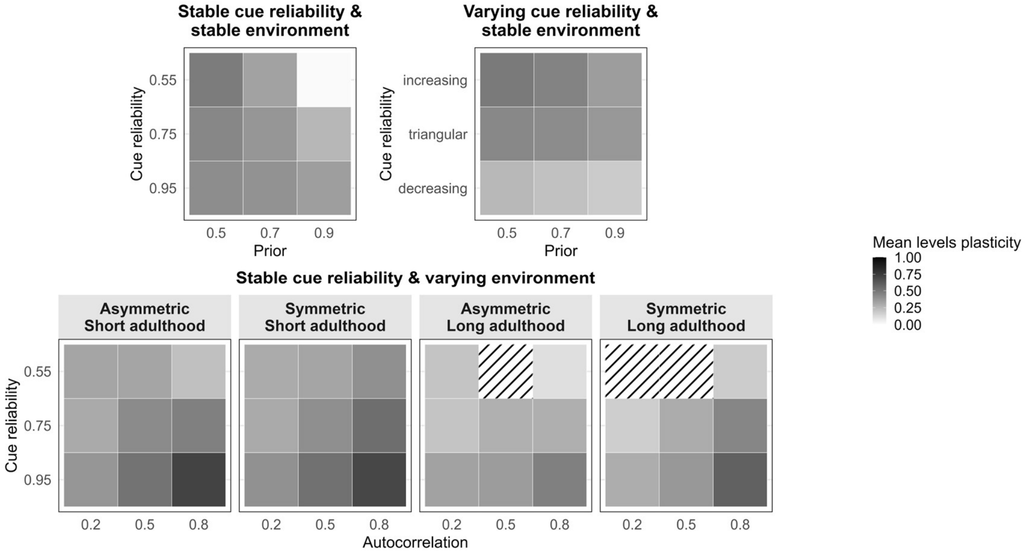


*Figure S14*: Average levels in plasticity. See, Figure S3 for the remainder of the legend.

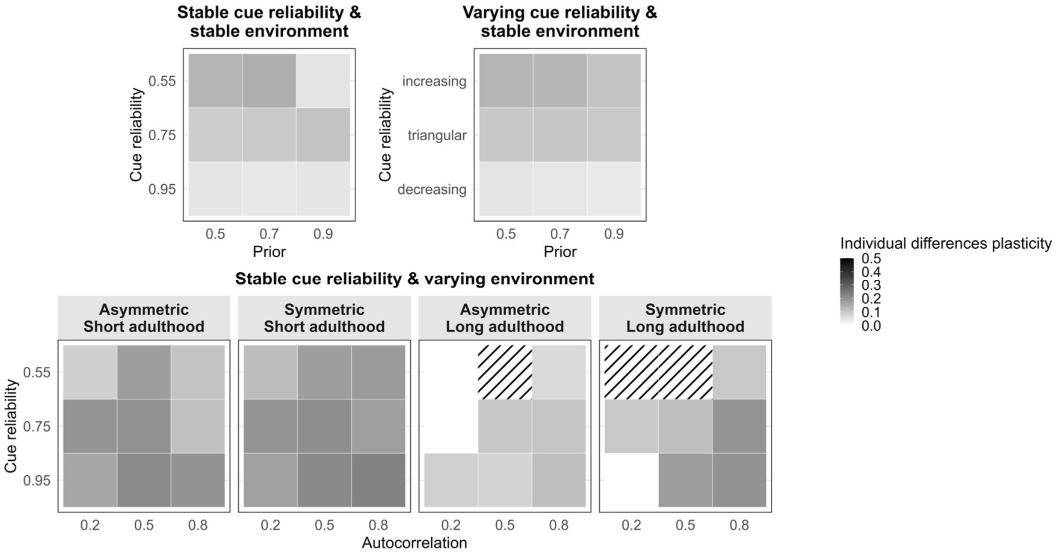


*Figure S15*: Individual differences in average levels of plasticity. See, Figure S3 for the remainder of the legend.

### Geometric-mean fitness values

**Table S1. Fitness comparison by strategy when the environment is stable within generations**

| prior | cue reliability | specialist | optimal policy | generalist | fitness |
|---|---|---|---|---|---|
| 0.5 | 0.55 | 11.00 | 12.82 | 11.00 | linear |
| 0.5 | 0.75 | 11.00 | 18.47 | 11.00 | linear |
| 0.5 | 0.95 | 11.00 | 20.72 | 11.00 | linear |
| 0.3 | 0.55 | 11.93 | 10.24 | 11.93 | linear |
| 0.3 | 0.75 | 11.93 | 18.81 | 11.93 | linear |
| 0.3 | 0.95 | 11.93 | 20.81 | 11.93 | linear |
| 0.1 | 0.55 | 15.80 | 15.49 | 15.80 | linear |
| 0.1 | 0.75 | 15.80 | 19.67 | 15.80 | linear |
| 0.1 | 0.95 | 15.80 | 20.85 | 15.80 | linear |
| 0.5 | 0.55 | 11.00 | 12.13 | 8.69 | increasing |
| 0.5 | 0.75 | 11.00 | 17.92 | 8.69 | increasing |
| 0.5 | 0.95 | 11.00 | 20.67 | 8.69 | increasing |
| 0.3 | 0.55 | 11.93 | 8.58 | 9.81 | increasing |

| prior | cue reliability | specialist | optimal policy | generalist | fitness |
|---|---|---|---|---|---|
| 0.3 | 0.75 | 11.93 | 18.22 | 9.81 | increasing |
| 0.3 | 0.95 | 11.93 | 20.72 | 9.81 | increasing |
| 0.1 | 0.55 | 15.80 | 15.49 | 14.61 | increasing |
| 0.1 | 0.75 | 15.80 | 19.20 | 14.61 | increasing |
| 0.1 | 0.95 | 15.80 | 20.78 | 14.61 | increasing |
| 0.5 | 0.55 | 11.00 | 14.23 | 13.31 | diminishing |
| 0.5 | 0.75 | 11.00 | 19.20 | 13.31 | diminishing |
| 0.5 | 0.95 | 11.00 | 20.87 | 13.31 | diminishing |
| 0.3 | 0.55 | 11.93 | 13.99 | 13.98 | diminishing |
| 0.3 | 0.75 | 11.93 | 19.37 | 13.98 | diminishing |
| 0.3 | 0.95 | 11.93 | 20.87 | 13.98 | diminishing |
| 0.1 | 0.55 | 15.80 | 15.74 | 16.80 | diminishing |
| 0.1 | 0.75 | 15.80 | 20.05 | 16.80 | diminishing |
| 0.1 | 0.95 | 15.80 | 20.91 | 16.80 | diminishing |

**Table S2. Fitness comparison by strategy when the environment changes within generations**

| symmArg | adultT | autocorr | cue reliability | specialist | optimal policy | generalist | fitness |
|---|---|---|---|---|---|---|---|
| symmetric | 1 | 0.8 | 0.55 | 11.00 | 11.91 | 11.00 | linear |
| symmetric | 1 | 0.8 | 0.75 | 11.00 | 14.61 | 11.00 | linear |
| symmetric | 1 | 0.8 | 0.95 | 11.00 | 15.02 | 11.00 | linear |
| symmetric | 1 | 0.5 | 0.55 | 11.00 | 11.20 | 11.00 | linear |
| symmetric | 1 | 0.5 | 0.75 | 11.00 | 12.00 | 11.00 | linear |
| symmetric | 1 | 0.5 | 0.95 | 11.00 | 12.80 | 11.00 | linear |
| symmetric | 1 | 0.2 | 0.55 | 11.00 | 11.12 | 11.00 | linear |
| symmetric | 1 | 0.2 | 0.75 | 11.00 | 11.62 | 11.00 | linear |
| symmetric | 1 | 0.2 | 0.95 | 11.00 | 12.12 | 11.00 | linear |
| symmetric | 20 | 0.8 | 0.55 | 11.00 | 11.22 | 11.00 | linear |
| symmetric | 20 | 0.8 | 0.75 | 11.00 | 11.89 | 11.00 | linear |

| symmArg | adultT | autocorr | cue reliability | specialist | optimal policy | generalist | fitness |
|---|---|---|---|---|---|---|---|
| symmetric | 20 | 0.8 | 0.95 | 11.00 | 11.99 | 11.00 | linear |
| symmetric | 20 | 0.5 | 0.55 | 11.00 | 11.02 | 11.00 | linear |
| symmetric | 20 | 0.5 | 0.75 | 11.00 | 11.10 | 11.00 | linear |
| symmetric | 20 | 0.5 | 0.95 | 11.00 | 11.18 | 11.00 | linear |
| symmetric | 20 | 0.2 | 0.55 | 11.00 | 11.01 | 11.00 | linear |
| symmetric | 20 | 0.2 | 0.75 | 11.00 | 11.04 | 11.00 | linear |
| symmetric | 20 | 0.2 | 0.95 | 11.00 | 11.07 | 11.00 | linear |
| asymmetric | 1 | 0.8 | 0.55 | 12.52 | 9.81 | 12.40 | linear |
| asymmetric | 1 | 0.8 | 0.75 | 12.52 | 12.93 | 12.40 | linear |
| asymmetric | 1 | 0.8 | 0.95 | 12.52 | 13.69 | 12.40 | linear |
| asymmetric | 1 | 0.5 | 0.55 | 11.22 | 6.21 | 11.22 | linear |
| asymmetric | 1 | 0.5 | 0.75 | 11.22 | 8.75 | 11.22 | linear |
| asymmetric | 1 | 0.5 | 0.95 | 11.22 | 9.49 | 11.22 | linear |
| asymmetric | 1 | 0.2 | 0.55 | 11.09 | 5.54 | 11.07 | linear |
| asymmetric | 1 | 0.2 | 0.75 | 11.09 | 7.22 | 11.07 | linear |
| asymmetric | 1 | 0.2 | 0.95 | 11.09 | 8.24 | 11.07 | linear |
| asymmetric | 20 | 0.8 | 0.55 | 13.45 | 15.84 | 12.97 | linear |
| asymmetric | 20 | 0.8 | 0.75 | 13.45 | 15.84 | 12.97 | linear |
| asymmetric | 20 | 0.8 | 0.95 | 13.45 | 15.84 | 12.97 | linear |
| asymmetric | 20 | 0.5 | 0.55 | 11.40 | 12.96 | 11.40 | linear |
| asymmetric | 20 | 0.5 | 0.75 | 11.40 | 12.96 | 11.40 | linear |
| asymmetric | 20 | 0.5 | 0.95 | 11.40 | 12.96 | 11.40 | linear |
| asymmetric | 20 | 0.2 | 0.55 | 11.16 | 12.23 | 11.25 | linear |
| asymmetric | 20 | 0.2 | 0.75 | 11.16 | 12.23 | 11.25 | linear |
| asymmetric | 20 | 0.2 | 0.95 | 11.16 | 12.23 | 11.25 | linear |
| symmetric | 1 | 0.8 | 0.55 | 11.00 | 12.00 | 8.69 | increasing |
| symmetric | 1 | 0.8 | 0.75 | 11.00 | 14.69 | 8.69 | increasing |
| symmetric | 1 | 0.8 | 0.95 | 11.00 | 15.10 | 8.69 | increasing |
| symmetric | 1 | 0.5 | 0.55 | 11.00 | 12.00 | 8.69 | increasing |

| symmArg | adultT | autocorr | cue reliability | specialist | optimal policy | generalist | fitness |
|---|---|---|---|---|---|---|---|
| symmetric | 1 | 0.5 | 0.75 | 11.00 | 15.17 | 8.69 | increasing |
| symmetric | 1 | 0.5 | 0.95 | 11.00 | 17.40 | 8.69 | increasing |
| symmetric | 1 | 0.2 | 0.55 | 11.00 | 12.00 | 8.69 | increasing |
| symmetric | 1 | 0.2 | 0.75 | 11.00 | 15.16 | 8.69 | increasing |
| symmetric | 1 | 0.2 | 0.95 | 11.00 | 18.56 | 8.69 | increasing |
| symmetric | 20 | 0.8 | 0.55 | 11.00 | 11.25 | 8.69 | increasing |
| symmetric | 20 | 0.8 | 0.75 | 11.00 | 12.24 | 8.69 | increasing |
| symmetric | 20 | 0.8 | 0.95 | 11.00 | 13.22 | 8.69 | increasing |
| symmetric | 20 | 0.5 | 0.55 | 11.00 | 11.10 | 8.69 | increasing |
| symmetric | 20 | 0.5 | 0.75 | 11.00 | 11.50 | 8.69 | increasing |
| symmetric | 20 | 0.5 | 0.95 | 11.00 | 11.90 | 8.69 | increasing |
| symmetric | 20 | 0.2 | 0.55 | 11.00 | 11.06 | 8.69 | increasing |
| symmetric | 20 | 0.2 | 0.75 | 11.00 | 11.31 | 8.69 | increasing |
| symmetric | 20 | 0.2 | 0.95 | 11.00 | 11.56 | 8.69 | increasing |
| asymmetric | 1 | 0.8 | 0.55 | 12.52 | 9.81 | 10.25 | increasing |
| asymmetric | 1 | 0.8 | 0.75 | 12.52 | 10.89 | 10.25 | increasing |
| asymmetric | 1 | 0.8 | 0.95 | 12.52 | 11.62 | 10.25 | increasing |
| asymmetric | 1 | 0.5 | 0.55 | 11.22 | 6.21 | 8.95 | increasing |
| asymmetric | 1 | 0.5 | 0.75 | 11.22 | 6.75 | 8.95 | increasing |
| asymmetric | 1 | 0.5 | 0.95 | 11.22 | 7.50 | 8.95 | increasing |
| asymmetric | 1 | 0.2 | 0.55 | 11.09 | 5.54 | 8.81 | increasing |
| asymmetric | 1 | 0.2 | 0.75 | 11.09 | 6.06 | 8.81 | increasing |
| asymmetric | 1 | 0.2 | 0.95 | 11.09 | 6.55 | 8.81 | increasing |
| asymmetric | 20 | 0.8 | 0.55 | 13.45 | 15.84 | 10.88 | increasing |
| asymmetric | 20 | 0.8 | 0.75 | 13.45 | 15.84 | 10.88 | increasing |
| asymmetric | 20 | 0.8 | 0.95 | 13.45 | 15.84 | 10.88 | increasing |
| asymmetric | 20 | 0.5 | 0.55 | 11.40 | 12.96 | 9.14 | increasing |
| asymmetric | 20 | 0.5 | 0.75 | 11.40 | 12.96 | 9.14 | increasing |
| asymmetric | 20 | 0.5 | 0.95 | 11.40 | 12.96 | 9.14 | increasing |

| symmArg | adultT | autocorr | cue reliability | specialist | optimal policy | generalist | fitness |
|---|---|---|---|---|---|---|---|
| asymmetric | 20 | 0.2 | 0.55 | 11.16 | 12.23 | 9.01 | increasing |
| asymmetric | 20 | 0.2 | 0.75 | 11.16 | 12.23 | 9.01 | increasing |
| asymmetric | 20 | 0.2 | 0.95 | 11.16 | 12.23 | 9.01 | increasing |
| symmetric | 1 | 0.8 | 0.55 | 11.00 | 13.44 | 13.31 | diminishing |
| symmetric | 1 | 0.8 | 0.75 | 11.00 | 14.65 | 13.31 | diminishing |
| symmetric | 1 | 0.8 | 0.95 | 11.00 | 14.66 | 13.31 | diminishing |
| symmetric | 1 | 0.5 | 0.55 | 11.00 | 13.31 | 13.31 | diminishing |
| symmetric | 1 | 0.5 | 0.75 | 11.00 | 13.25 | 13.31 | diminishing |
| symmetric | 1 | 0.5 | 0.95 | 11.00 | 13.09 | 13.31 | diminishing |
| symmetric | 1 | 0.2 | 0.55 | 11.00 | 13.33 | 13.31 | diminishing |
| symmetric | 1 | 0.2 | 0.75 | 11.00 | 13.31 | 13.31 | diminishing |
| symmetric | 1 | 0.2 | 0.95 | 11.00 | 13.21 | 13.31 | diminishing |
| symmetric | 20 | 0.8 | 0.55 | 11.00 | 13.36 | 13.31 | diminishing |
| symmetric | 20 | 0.8 | 0.75 | 11.00 | 13.36 | 13.31 | diminishing |
| symmetric | 20 | 0.8 | 0.95 | 11.00 | 13.26 | 13.31 | diminishing |
| symmetric | 20 | 0.5 | 0.55 | 11.00 | 13.37 | 13.31 | diminishing |
| symmetric | 20 | 0.5 | 0.75 | 11.00 | 13.33 | 13.31 | diminishing |
| symmetric | 20 | 0.5 | 0.95 | 11.00 | 13.30 | 13.31 | diminishing |
| symmetric | 20 | 0.2 | 0.55 | 11.00 | 13.37 | 13.31 | diminishing |
| symmetric | 20 | 0.2 | 0.75 | 11.00 | 13.35 | 13.31 | diminishing |
| symmetric | 20 | 0.2 | 0.95 | 11.00 | 13.34 | 13.31 | diminishing |
| asymmetric | 1 | 0.8 | 0.55 | 12.52 | 11.85 | 14.40 | diminishing |
| asymmetric | 1 | 0.8 | 0.75 | 12.52 | 14.95 | 14.40 | diminishing |
| asymmetric | 1 | 0.8 | 0.95 | 12.52 | 15.59 | 14.40 | diminishing |
| asymmetric | 1 | 0.5 | 0.55 | 11.22 | 13.03 | 13.47 | diminishing |
| asymmetric | 1 | 0.5 | 0.75 | 11.22 | 12.88 | 13.47 | diminishing |
| asymmetric | 1 | 0.5 | 0.95 | 11.22 | 12.77 | 13.47 | diminishing |
| asymmetric | 1 | 0.2 | 0.55 | 11.09 | 13.24 | 13.33 | diminishing |
| asymmetric | 1 | 0.2 | 0.75 | 11.09 | 13.29 | 13.33 | diminishing |

| symmArg | adultT | autocorr | cue reliability | specialist | optimal policy | generalist | fitness |
|---|---|---|---|---|---|---|---|
| asymmetric | 1 | 0.2 | 0.95 | 11.09 | 13.07 | 13.33 | diminishing |
| asymmetric | 20 | 0.8 | 0.55 | 13.45 | 15.85 | 14.82 | diminishing |
| asymmetric | 20 | 0.8 | 0.75 | 13.45 | 15.85 | 14.82 | diminishing |
| asymmetric | 20 | 0.8 | 0.95 | 13.45 | 15.80 | 14.82 | diminishing |
| asymmetric | 20 | 0.5 | 0.55 | 11.40 | 13.77 | 13.60 | diminishing |
| asymmetric | 20 | 0.5 | 0.75 | 11.40 | 13.74 | 13.60 | diminishing |
| asymmetric | 20 | 0.5 | 0.95 | 11.40 | 13.73 | 13.60 | diminishing |
| asymmetric | 20 | 0.2 | 0.55 | 11.16 | 13.52 | 13.46 | diminishing |
| asymmetric | 20 | 0.2 | 0.75 | 11.16 | 13.52 | 13.46 | diminishing |
| asymmetric | 20 | 0.2 | 0.95 | 11.16 | 13.48 | 13.46 | diminishing |
